# Multimodal Imaging and Logistic Weighted Cognitive Scores for Classification of MCI, AD, and FTD Subtypes

**DOI:** 10.1101/2025.09.27.678895

**Authors:** Sunil Kumar Khokhar, Rohit Misra, Manoj Kumar, Faheem Arshad, Nithin Thanissery, Sandeep Kumar, Tejaswini Manae, Subasree Ramakrishnan, Sandhya Mangalore, Suvarna Alladi, Tapan Kumar Gandhi, Rose Dawn Bharath

## Abstract

**Background:** Differentiating between mild cognitive impairment (MCI), Alzheimer’s disease (AD), and frontotemporal dementia (FTD) subtypes remains a clinical challenge due to overlapping cognitive symptoms, structural atrophy, and metabolic patterns, especially in the early stages. Multimodal classification approaches integrating neuroimaging and cognitive scores may offer early and accurate characterization and subsequently improved diagnostic precision.

**Methods:** In this study, we included 100 participants (50 AD, 30 FTD, including 14 bvFTD and 16 PPA, and 20 MCI) who underwent simultaneous structural MRI and FDG-PET imaging. Cortical thickness (CTH) from anatomical MRI and standardized uptake values from FDG-PET were extracted using FreeSurfer and PETSurfer pipelines, respectively. CTH and FDG-PET features were combined into a single vector through a logistic weighting function derived from ACE-III scores, capturing the progressive nature of cognitive decline in dementia. A Naive Bayes classifier was then trained to differentiate diagnostic groups based on the merged features.

**Results:** The model achieved classification accuracies of 83% for MCI vs. dementia (AD + FTD), 85% for MCI vs. FTD, 87% for MCI vs. PPA, 71% for MCI vs. bvFTD, 64% for MCI vs. AD, and 69% for AD vs. FTD. The overall classification accuracy was 68%, with the highest discriminative performance observed in separating MCI from FTD subtypes.

**Conclusions:** This study presents a novel, cognition-weighted multimodal approach combining structural and metabolic imaging to enhance the classification of neurodegenerative syndromes. Findings from this study, underscore the potential of integrating ACE-III scores with neuroimaging biomarkers for accurate characterization and, early-stage differentiation of MCI, AD, and FTD variants.

## INTRODUCTION

Alzheimer’s Disease (AD) and Frontotemporal Dementia (FTD) represent the most prevalent neurodegenerative diseases contributing to cognitive and behaviour impairment. FTD presents clinically as variants including the behavioral variant FTD (bvFTD) with behavioral and executive dysfunctions (Rascovsky et al., 2011a), and primary progressive aphasia (PPA) encompassing the semantic variant PPA (svPPA) characterized by semantic anomalies and comprehension deficits, and non-fluent/agrammatic variant PPA (nfvPPA), impairing speech, grammar, and word production (Gorno-Tempini et al., 2011), AD characterized by progressive memory loss and contributes significantly to cognitive impairment. Prompt detection and accurate identification of these neurodegenerative conditions, and their prodromal phase, mild cognitive impairment (MCI), (Albert et al., 2011; Petersen et al., 2009) are critical for early diagnosis.

Neuroimaging tools like magnetic resonance imaging (MRI) and [18-F] Fluorodeoxyglucose Positron emission tomography (FDG-PET) studies provide adjunctive value in the diagnosis of these disorders, although interpretation largely depends on the expertise of the reporting clinician (Finger, 2016; Foster et al., 2007). Novel advances in the quantitative neuroimaging field permit the automated study of cortical morphometry and cerebral metabolic alterations. Previous studies revealed medial temporal and parietotemporal atrophy in AD, while FTD syndromes exhibit atrophy in the frontal, anterior temporal, and fronto-insular cortices (Gorno-Tempini et al., 2011; Herholz et al., 2011; Ou et al., 2019; Qiao et al., 2022). For instance, bvFTD typically exhibits pronounced atrophy in the anterior cingulate and frontal insular cortices, whereas nfv-PPA presents atrophy predominantly in the left frontal region, including the inferior frontal gyrus, pars triangularis, rolandic operculum, and precentral gyrus. svPPA is characterized by distinctive atrophy primarily in the left anterior temporal lobe, making it distinguishable from other FTD syndromes (Gorno-Tempini et al., 2011). However, delineating these different dementia types presents a challenge due to the subtlety of early-stage FTD cortical atrophy and overlapping atrophy characteristics across FTD syndromes, AD, and MCI. Machine learning (ML) algorithms have gained momentum for the classification and prediction of diverse neurodegenerative disorders (Cuingnet et al., 2011; Ito et al., 2015).

The previous literature of neurodegenerative disease classification, particularly concerning AD and FTD, has witnessed significant advancements through the integration of imaging biomarkers and cortical thickness (CTH) measures. One notable study explored the effectiveness of cerebrospinal fluid (CSF) and MRI biomarkers across a spectrum of neurodegenerative diseases. It highlighted significant variances in imaging variables such as hippocampal and amygdalar volumes, and CTH across different clinical diagnoses, affirming the utility of these biomarkers in clinical practice beyond AD and MCI groups (Kaipainen et al., 2020). In another study, a hierarchical classification model of FTD was developed using ML approaches to assess the CTH. The research utilized the Small Univalue Segment Assimilating Nucleus (SUSAN) technique to minimize noise, enhancing the clarity and reliability of the imaging data. This model achieved significant accuracies, with the best performance noted in the support vector machine (SVM) approach, illustrating the potential of ML-driven methods in improving the diagnostic predictability of neurodegenerative diseases (Poonam et al., 2021). Furthermore, a study on ML-based hierarchical classification demonstrated the potential of CTH data to differentiate between FTD, AD, and their subtypes (Kim et al., 2019a). The use of noise-reducing techniques and a structured hierarchical classification approach yielded an overall accuracy of 75.8%; the classification accuracies of CN vs Dementia, AD vs FTD, and bvFTD vs PPA were 86.1%, 90.8%, and 86.9%, respectively (Kim et al., 2019a). This approach not only enhanced the accuracy of classification but also provided critical insights into the specific regional patterns of neurodegeneration, paving the way for targeted interventions (Kim et al., 2019b). Other studies in the classification of AD patients from CN subjects showed accuracy of 92.35% and 91.50 %, and MCI from AD with the accuracy of 75.90% (Wee et al., 2012; Westman et al., 2012). The current investigation utilised a unique method using a logistic function to assign weights derived from cognitive scores, specifically the Addenbrooke’s Cognitive Examination-III (ACE-III) score, to the CTH and FDG-PET variables. The performance of the classification models is evaluated through various metrics, including the accuracy, sensitivity, specificity, and AUC-ROC. The current findings indicate that a Naive Bayes classifier with PCA for feature extraction provides promising outcomes on unified CTH and FDG-PET datasets, although with varying accuracy levels according to severity. These results underscore the promising potential of these ML algorithms in the early diagnosis and more accurate classification of MCI, AD, bvFTD, and PPA.

## METHODS

### Participants

In this ambispective comparative cohort study, we investigated a total of one hundred patients with dementia, comprising of 50 individuals with AD, 20 with MCI, and 30 with FTD. The FTD group had two subgroups: 14 with the bvFTD and 16 with PPA. These patients were selected consecutively from the patients referred for MRI-PET investigation for their routine clinical care. The severity of dementia was assessed using the Clinical Dementia Rating (CDR) scale. Cognitive evaluations were conducted using ACE-III (Mekala et al., 2020), which encompassed domains such as memory, attention, language, fluency, and visuospatial skills. The diagnosis of FTD was made in patients using the standard criteria (Gorno-Tempini et al., 2011; Rascovsky et al., 2011b), and AD was diagnosed in patients who fulfilled the NIA-AA (National Institute on Aging and Alzheimer’s Association) criteria for probable and possible AD (McKhann et al., 2011), while the diagnosis of MCI was made based on the modified Petersen criteria (Petersen, 2004). This study received approval from the institute’s ethics committee [Ethics NO. NIMH/DO/(BS&NS DIV.) 2019-20].

There were no significant between-group differences with respect to age, duration of illness, and education level (p>0.05). However, differences were observed in the ACE scores and its subdomain scores among the groups. The mean ACE scores for MCI, AD, FTD-BV, and PPA were 87.3 (SD=5.53), 46.02 (SD=20.61), 52.57 (SD=18.79), and 33.56 (SD=21.09), respectively. These differences were statistically significant (p<0.001). Post hoc Bonferroni tests (Supplementary Table 1) showed that scores in the MCI group were significantly higher than those of all other groups (p<0.01), additionally bvFTD showed significantly higher ACE with subdomain of language scores in compare to PPA, while fluency and language score were significantly lower in PPA in comparison to AD (Table 1). The participants in this study were the same cohort as those included in the previously reported AD and FTD studies (Khokhar et al., 2023, 2025), and thus, the acquisition protocols and behavioral assessments were consistent across all studies.

**Table 1:**
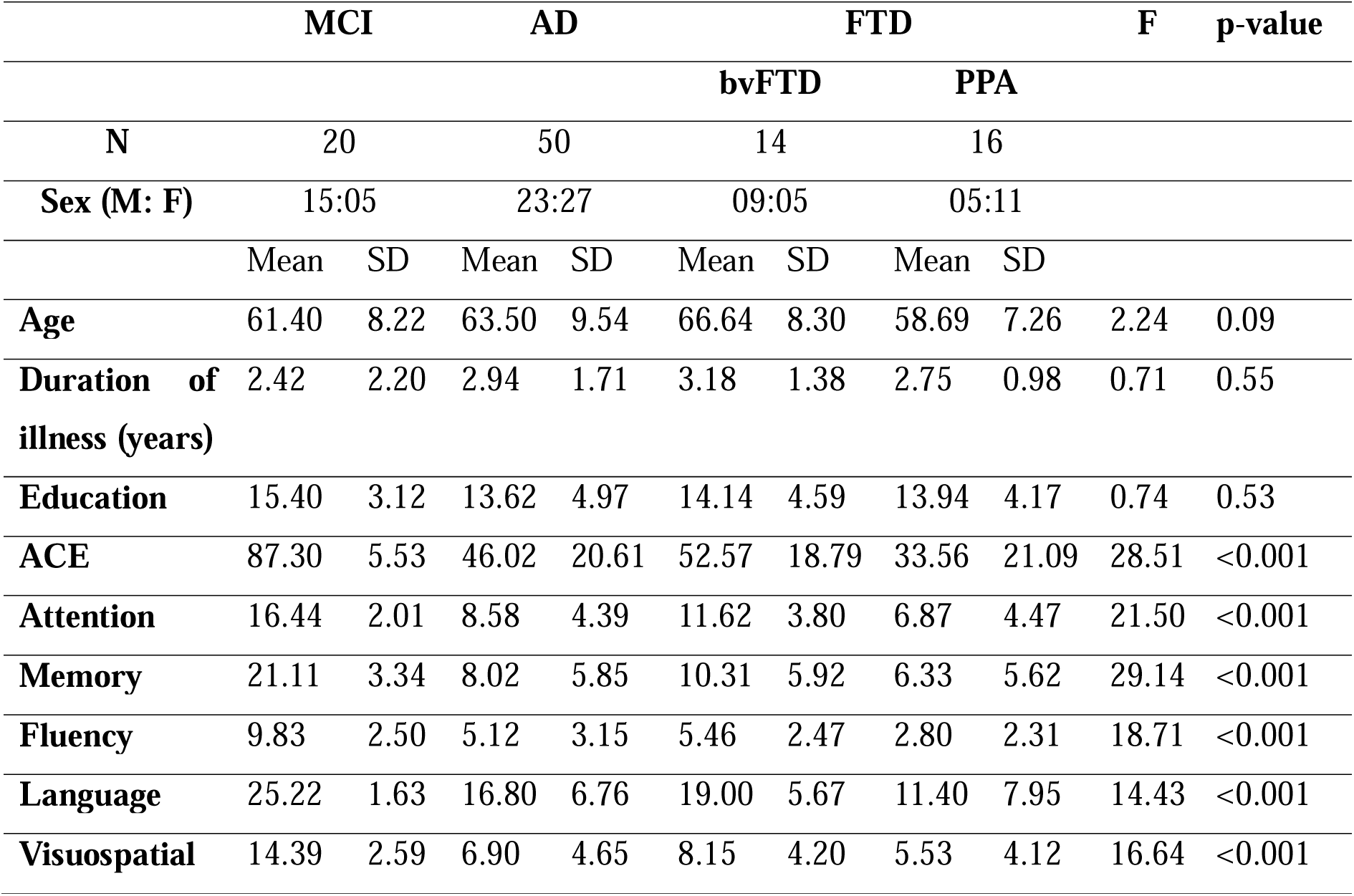
Demographic Information of Participants.

### Acquisition parameters

#### MRI

All participants underwent structural magnetic resonance imaging (sMRI) using a 3 Tesla clinical MRI (Biograph mMR) scanner (Siemens, Erlangen, Germany) equipped with a 16-channel head coil. To minimize head movement and enhance patient comfort during the scan, the head was stabilized with foam pads. The imaging process included a 3D T1-weighted Magnetization Prepared Rapid Acquisition Gradient-Recalled Echo (MPRAGE) sequence with 192 sections and a repetition time (TR) of 2300ms, echo time (TE) of 2.42 ms, slice thickness: 1 mm, flip angle of 9°, a field of view (FOV) of 250 x 250mm, a matrix resolution of 256 x 256. In addition, other sequences such as Fluid-Attenuated Inversion Recovery (FLAIR), T2-weighted imaging, and Susceptibility Weighted Imaging (SWI) were employed as a routine diagnostic MRI scan and also ensuring the exclusion of structural abnormalities in all patients.

#### FDG-PET

All participants received FDG 185± 10%, MBq (5.0 mCi) 30 minutes prior to image acquisition, and imaging was performed in the list mode for 15 min. These images had a resolution of 256 matrices, a 300mm field of view, and voxel dimensions of 2.3mm×2.3mm×5mm. using fully 3D or Fourier-rebinned Ordered Subset Expectation Maximization (OSEM) iterative reconstruction algorithms with three iterations and 35 subsets. Standard corrections for attenuation, scatter, random coincidences, and decay were applied, as well as a 5 mm Gaussian post-reconstruction filter.

### Image Processing

#### Cortical Thickness

The cortical reconstruction and estimation of CTH were performed using the FreeSurfer image analysis tool v6.05 (Fischl, 2012) (https://surfer.nmr.mgh.harvard.edu). The pipeline within the FreeSurfer suite involves the automated procedure for motion correction, skull stripping, automated transformation onto Talairach space, subcortical white matter and gray matter segmentation, and tessellation of gray matter and white matter boundaries. Next, the cortical surface was reconstructed by high-resolution inter-subject alignment procedures to yield a high-resolution quantification of CTH, along with differentiation into different atlas-derived and prescribed cortical brain regions (Dale et al., 1999). The Desikan Killiany (DK) atlas was used for the parcellation of the cortical surface into distinct units based on gyral and sulcal anatomy, with 34 regions per hemisphere (Desikan et al., 2006).

#### FDG-PET

The FDG-PET image processing was performed using the PETSurfer analysis tool v6.05 (Greve et al., 2014). The process started by capitalizing on the FreeSurfer results from the MR T1 image data to enable PET processing. This involved registration into a standardized template space and employing the Partial Volume Correction (PVC) using geometric transfer matrix (GTM) techniques for partial volume effects (Greve et al., 2016). After the segmentation and registration processes, PETSurfer was able to compute the SUV values using the pons as a reference area, calculate the volume size in the region, and measure voxel variance. We then used a 10mm full-width half maximum (FWHM) Gaussian filter to smooth the FDG-PET images. We also applied the DK atlas to parcel the cortical surface into 34 regions per hemisphere (Desikan et al., 2006).

**Figure 1:**
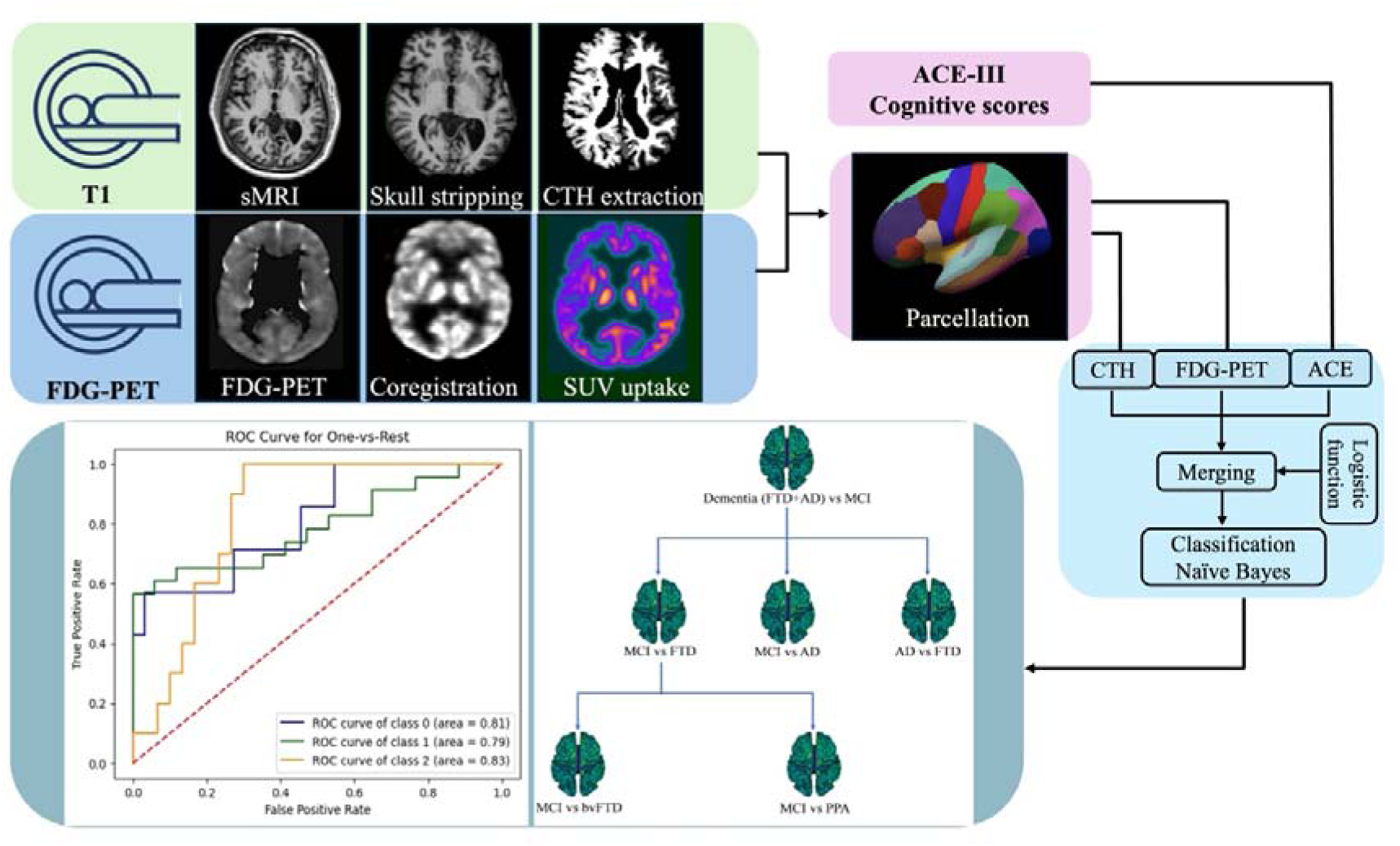
Demonstrating integration of multimodal MRI and FDG-PET processing and classification pipeline. T1 and FDG-PET images were processed for CTH and SUV extraction, respectively. ACE-III cognitive scores and parcellated regions were integrated with imaging data using a logistic weighting function. A Naive Bayes model classified MCI, AD, and FTD, with performance shown via ROC curves and a decision tree.

### Machine learning-based classifications

We used the scikit-learn library for ML classification in Jupiter notebook (Python v3.7.3), to conduct ML classification algorithms (Pedregosa et al., 2011). Here, we used a Naive Bayes model for the classification of MCI, AD, and FTD using CTH, FDG-PET, and cognitive score.

#### Naive Bayes

Naive Bayes is a supervised ML algorithm that is based on certain “naive” assumptions that predictors in a model are independent of each other. Though unrealistic in a real-world scenario, this simplistic model makes the calculation of probabilities much simpler and enables the algorithm to be fast and efficient. Despite this simplifying assumption, Naive Bayes often provides good results, especially for large datasets with many features.

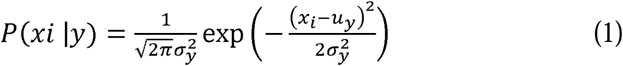

P(*xi* |*y*) - represents the probability of a feature *x_i_* given class of *y*

*u_y_* - is the mean of feature *x_i_ for class y*

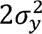 - is the variance of feature *x_i_ for class y*

The term 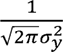 is the normalization factor of the Gaussian distribution. The exponential term 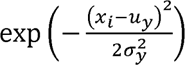 calculates the probability density of *x_i_* for a Gaussian distribution with a mean *u_y_* and variance 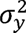. The parameters, *σ_y_* and *u_y_* are estimated using maximum likelihood.

### Merging process

It was observed that the correlation coefficient of PET SUV values were significantly higher than the CTH values. Hence, we used ACE scores to normalise both data sets. Merging datasets was feasible as it was acquired simultaneously in a simultaneous MR-PET scanner. Weight’s function was defined to accept an input from ACE scores and compute the weights CTH and FDG-PET using a logistic function. The logistic function was applied to each element of the ACE scores to calculate the corresponding FDG-PET values. The CTH values were calculated as 1 minus the FDG-PET values. The CTH and FDG-PET weights were calculated for each element of the ACE scores using the weight function. A merged array was initialized with the same shape as data CTH. The algorithm iterated through each region of interest (ROI) from 0 to 67. For each ROI of CTH was standardized by subtracting the mean and dividing by the standard deviation, while FDG-PET ROIs were also standardized in the same manner. The resulting products were added together to create the merged data for each ROIs. Finally merged two sets of data, CTH and FDG-PET, using the weights CTH and FDG-PET derived from ACE.

### Logistic function

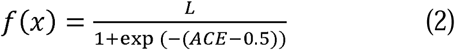

ACE score was normalised for 0 to 1.

*L* = the curve’s maximum value

### Statistical analysis

Statistical analysis was performed with ‘*scipy’* library in Python. Differences between groups in age, education, and ACE-III were tested using an analysis of variance (ANOVA) for continuous variables, while gender differences between groups were tested using a chi-square test. Post-hoc analyses using Bonferroni-corrected pairwise t-tests, following a Kruskal-Wallis test for the ordinal variable CDR, were conducted to examine pairwise group differences with a significance level of 0.05.

## RESULTS

The Naive Bayes classifier was implemented for the classification of three different classes of dementia: MCI, AD, and FTD. The obtained accuracy for the model was 0.68, suggesting that in 68% of instances, the model could correctly classify these conditions (Table 2). The area under the receiver operating characteristic curve (AUC-ROC) score for AD was 0.79, for MCI was 0.81, and for FTD was 0.83. The sensitivity (the true positive rate) for AD, MCI, and FTD was 0.57, 0.78, and 0.50, respectively, while the specificity (true negative rate) for these classes was 0.84, 0.77, and 0.87, respectively (Table 3).

**Figure 2:**
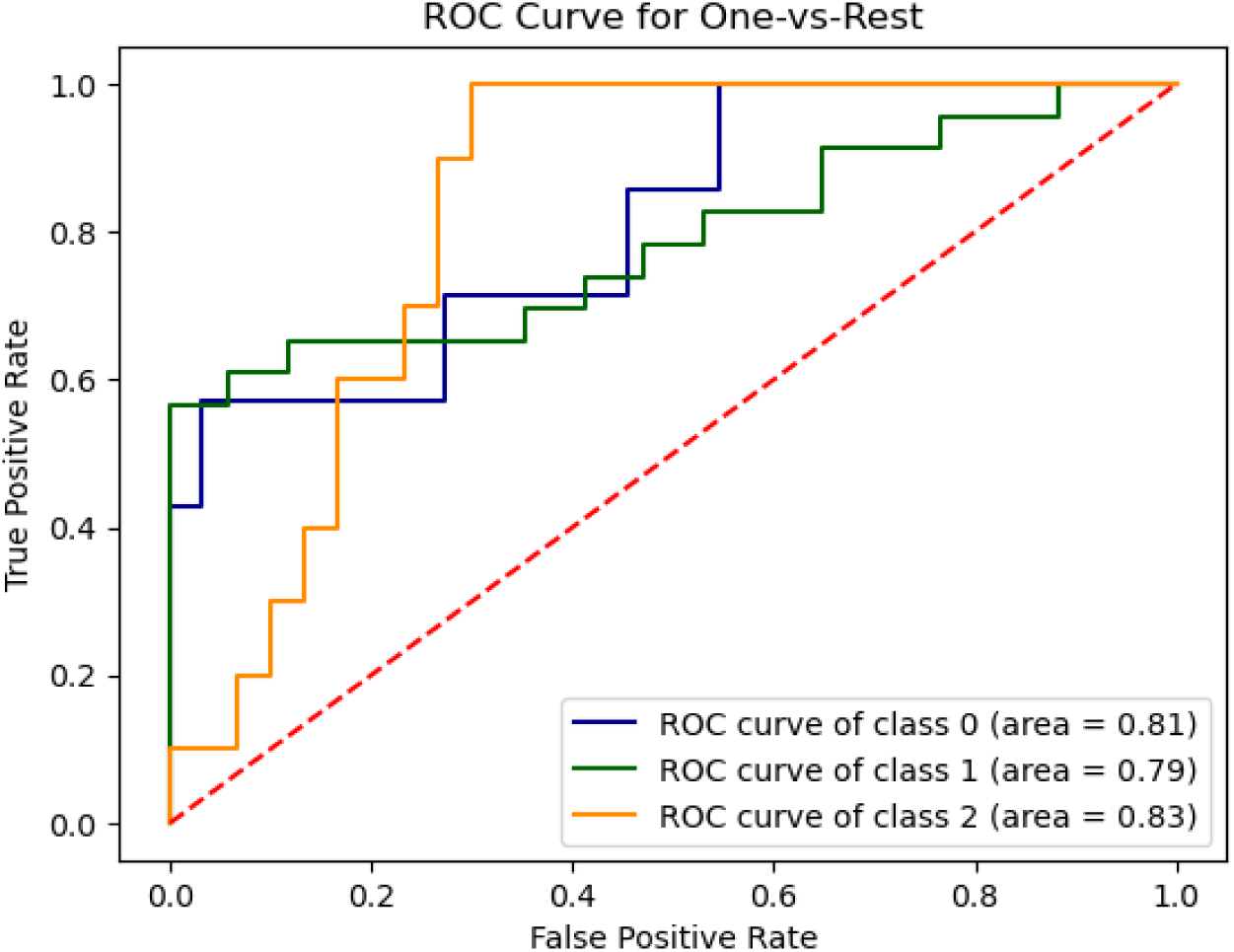
The figure displays the ROC curve for the One-vs-Rest approach.

**Figure 3:**
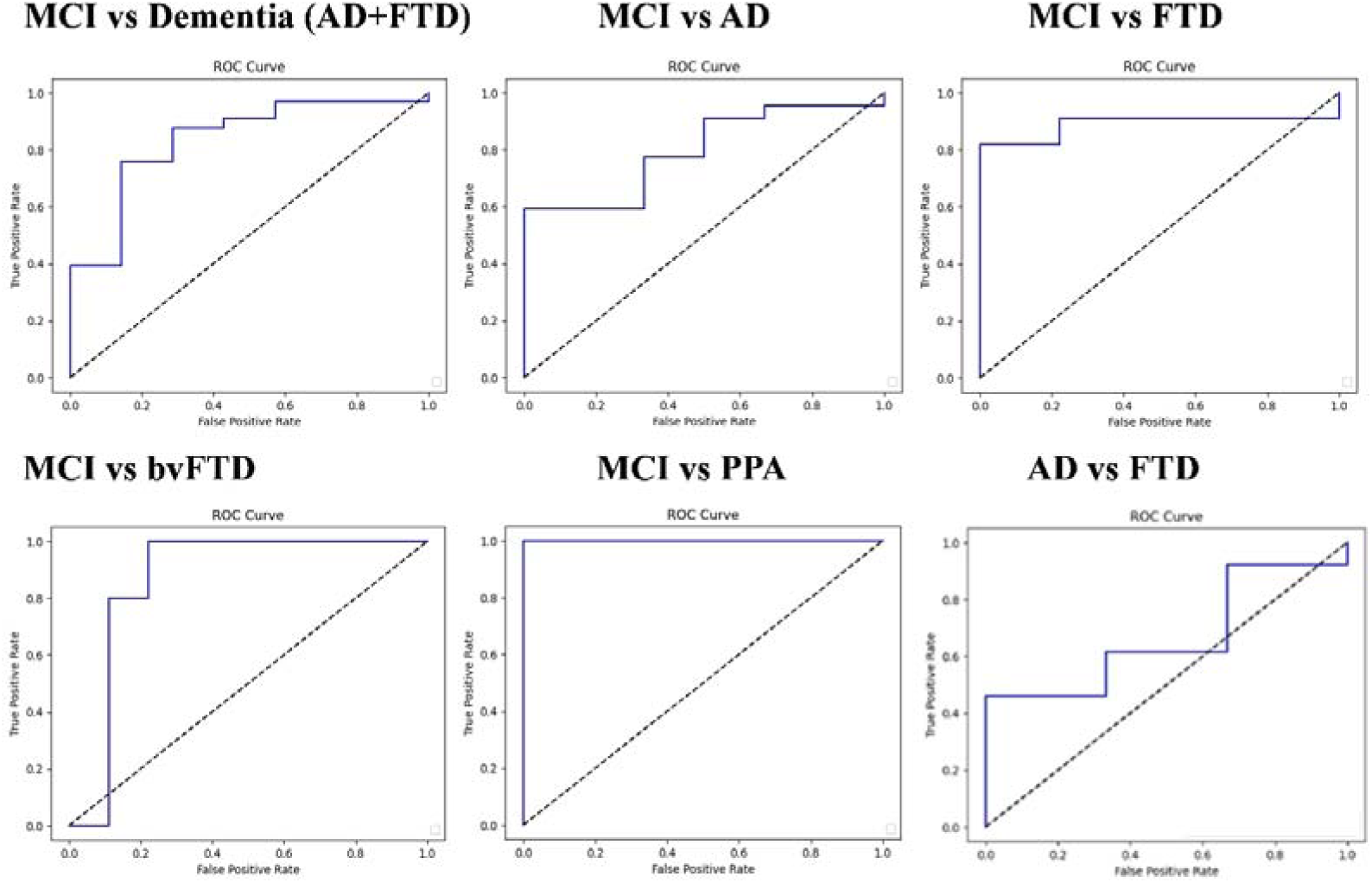
ROC curves depicting the discriminative ability of the Naive Bayes classifier in differentiating MCI from various dementia types (AD, FTD, bvFTD, PPA).

**Table 2:** The table exhibits the performance metrics of the Naive Bayes classifier for the different classification tasks, offering a unified view of its effectiveness.

| Metric | MCI | AD | FTD |
| --- | --- | --- | --- |
| Accuracy | 0.68 |  |  |
| AUC-ROC score | 0.81 | 0.79 | 0.83 |
| Sensitivity | 0.78 | 0.57 | 0.50 |
| Specificity | 0.77 | 0.84 | 0.87 |

**Table 3:**
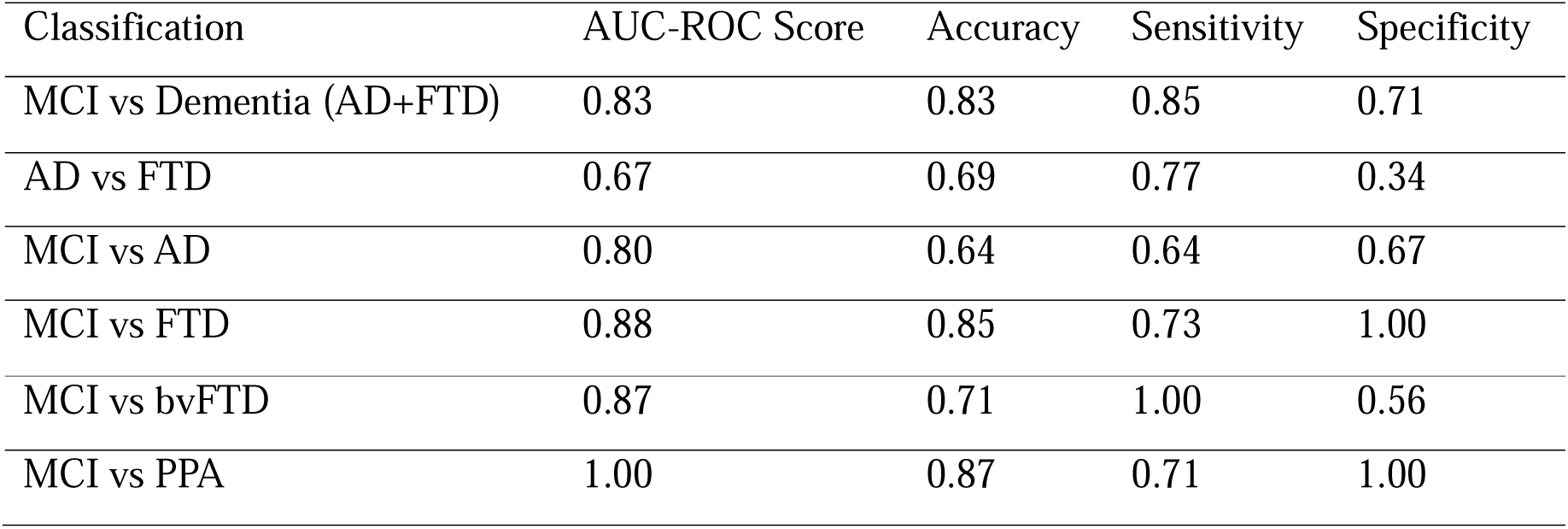
The table exhibits the performance metrics of the Naive Bayes classifier for the different classification tasks, offering a unified view of its effectiveness.

### Performance metrics for Naive Bayes classifier (MCI, AD, FTD)

Subsequently, the Naive Bayes classifier was evaluated on its ability to distinguish MCI from Dementia (AD and FTD combined), MCI from AD, MCI from FTD, MCI from bvFTD, and MCI from PPA. The AUC-ROC scores, accuracy, sensitivity, and specificity for each classification task are provided below.

### Performance metrics for Naive Bayes classifier

When distinguishing between MCI and a combined group of AD and FTD, the classifier had an AUC-ROC score of 0.83, an accuracy of 0.83, a sensitivity of 0.85, and a specificity of 0.71. This suggests that the classifier is quite capable of differentiating between these conditions. In the task of differentiating MCI from AD specifically, the classifier had an AUC-ROC score of 0.80, an accuracy of 0.64, a sensitivity of 0.64, and a specificity of 0.67. The model was reasonably good at identifying and differentiating these conditions, though not as proficient as with the combined group. The classifier showed a stronger performance when differentiating between MCI and FTD, with an AUC-ROC score of 0.88, accuracy of 0.85, sensitivity of 0.73, and specificity of 1.0. This indicates a relatively good ability to discriminate between these two conditions. Differentiating MCI from bvFTD saw the classifier achieve an AUC-ROC score of 0.87, accuracy of 0.71, a perfect sensitivity score of 1.0, and specificity of 0.56. Despite the lower specificity, the results collectively suggest an effective discrimination between MCI and bvFTD. Finally, when the task was to differentiate MCI from PPA, the classifier excelled with a perfect AUC-ROC score of 1.0, accuracy of 0.87, sensitivity of 0.71, and a perfect specificity of 1.0. This indicates the classifier is highly capable of distinguishing between these two conditions.

## DISCUSSION

In this study, we introduce a novel approach for classifying dementia subtypes by integrating CTH from structural MRI, SUV uptake values from FDG-PET, and cognitive assessment using ACE-III, further using a Naive Bayes classifier combined with a logistic function-based weighting function. Our results underscore the diagnostic utility of combining structural, metabolic, and cognitive biomarkers, particularly in differentiating MCI from AD and FTD subtypes. The classifier achieved the highest accuracy in differentiating MCI from PPA (87%) and FTD overall (85%), and 83% accuracy when distinguishing MCI from all dementia groups combined, AD & FTD. These findings align with prior evidence that FDG-PET provides greater sensitivity for early metabolic dysfunction, while cortical atrophy measured by CTH adds spatial specificity to later-stage degeneration (Herholz & Ebmeier, 2011; Jack Jr. et al., 2018). The hybrid model’s performance supports recent trends in multimodal neuroimaging that emphasize the added value of combining structural and functional data for early differential diagnosis (Bi et al., 2024; Khokhar et al., 2025; Venugopalan et al., 2021).

Cognitive assessments remain clinically crucial for diagnosing and staging dementia. The ACE-III is a validated tool encompassing attention, memory, fluency, language, and visuospatial domains. Our approach leverages ACE-III scores not merely as a covariate, but as a dynamic weight through a logistic function, modulating the influence of imaging features in classification. This method models the non-linear trajectory of neurodegeneration and reflects growing evidence that cognitive decline correlates with the convergence of multiple neuropathological processes (Lawry Aguila et al., 2025; Marinescu et al., 2019). Unlike traditional models with static or uniform feature weighting, our biologically motivated framework captures the progressive and heterogeneous nature of dementia. ML approaches are increasingly used in dementia research to improve diagnostic accuracy (Wang et al., 2024). While support vector machines, random forests, and neural networks have shown promise, they often require large datasets and extensive tuning. Naive Bayes classifiers, on the other hand, are computationally simple and provide transparent probabilistic outputs. Their assumption of feature independence, although idealized, has been shown to work effectively in neuroimaging classification tasks where sample sizes are limited (Moradi et al., 2015; Bansal et al., 2020). Our results confirm that Naive Bayes performs robustly in high-dimensional, low-sample contexts and is well-suited for translational research.

A central contribution of this study is the use of a logistic function applied to ACE-III scores to derive weights for imaging features. This biologically motivated approach reflects the disease continuum, where declining cognitive performance is increasingly influenced by metabolic deficits, thus enhancing the physiological relevance of data fusion. Unlike traditional multimodal models that often rely on equal or unsupervised feature weighting (Sadiq et al., 2022; Venugopalan et al., 2021), our method provides a flexible and scalable mechanism to integrate cognition-adjusted biomarkers. The use of a Naive Bayes classifier, though less common than support vector machines and deep learning models, proved efficient and interpretable. Despite its simplifying assumption of feature independence, it offered robust performance in a high-dimensional, small-sample context, consistent with earlier studies demonstrating its effectiveness in related domains (Kubi & Nazir, 2025; Moradi et al., 2015; Zhang et al., 2011). Notably, the classifier’s specificity in distinguishing MCI from both bvFTD and PPA reached 100%, indicating a strong potential for ruling out false positives in early clinical screening.

However, moderate sensitivity in distinguishing AD (57%) likely reflects the heterogeneity within the AD cohort, including early-stage or atypical presentations that blur boundaries with other dementias (Vogel et al., 2021). The overlap in structural and metabolic patterns between AD and FTD subtypes, especially in the frontal cortex, poses a well-known diagnostic challenge (Musa et al., 2020). Future models may benefit from ensemble classifiers or non-linear approaches such as gradient boosting and neural networks that better capture these complex overlaps. Despite its strengths, this study has limitations. The sample size, particularly for FTD subtypes, was modest, and although results are promising, larger, multi-center validation is needed. Additionally, while ACE-III scores were effectively used to derive weights, they may be influenced by demographic factors such as education or language fluency. Expanding the weighting model to include alternate cognitive or functional scores (e.g., MoCA, MMSE, and digital cognitive testing) could improve generalizability. Moreover, while simultaneous MR-PET acquisition facilitated data fusion in this study, many clinical settings rely on asynchronous scans. Developing preprocessing pipelines to account for temporal variability will be essential for clinical translation. Finally, longitudinal designs assessing diagnostic stability over time would strengthen the clinical utility of this framework and allow prediction of disease progression.

### Conclusion

This study highlights the potential of physiologically informed, cognition-weighted integration of multimodal imaging for the classification of dementia syndromes. The approach leverages structural and metabolic features aligned with cognitive severity, providing a more nuanced diagnostic model. Future work should explore generalization across larger cohorts and integrate additional biomarkers, such as CSF tau/Aβ, plasma NfL, or digital phenotyping, to further refine early diagnosis and personalized care pathways.

---

**Supplementary Table 1:**
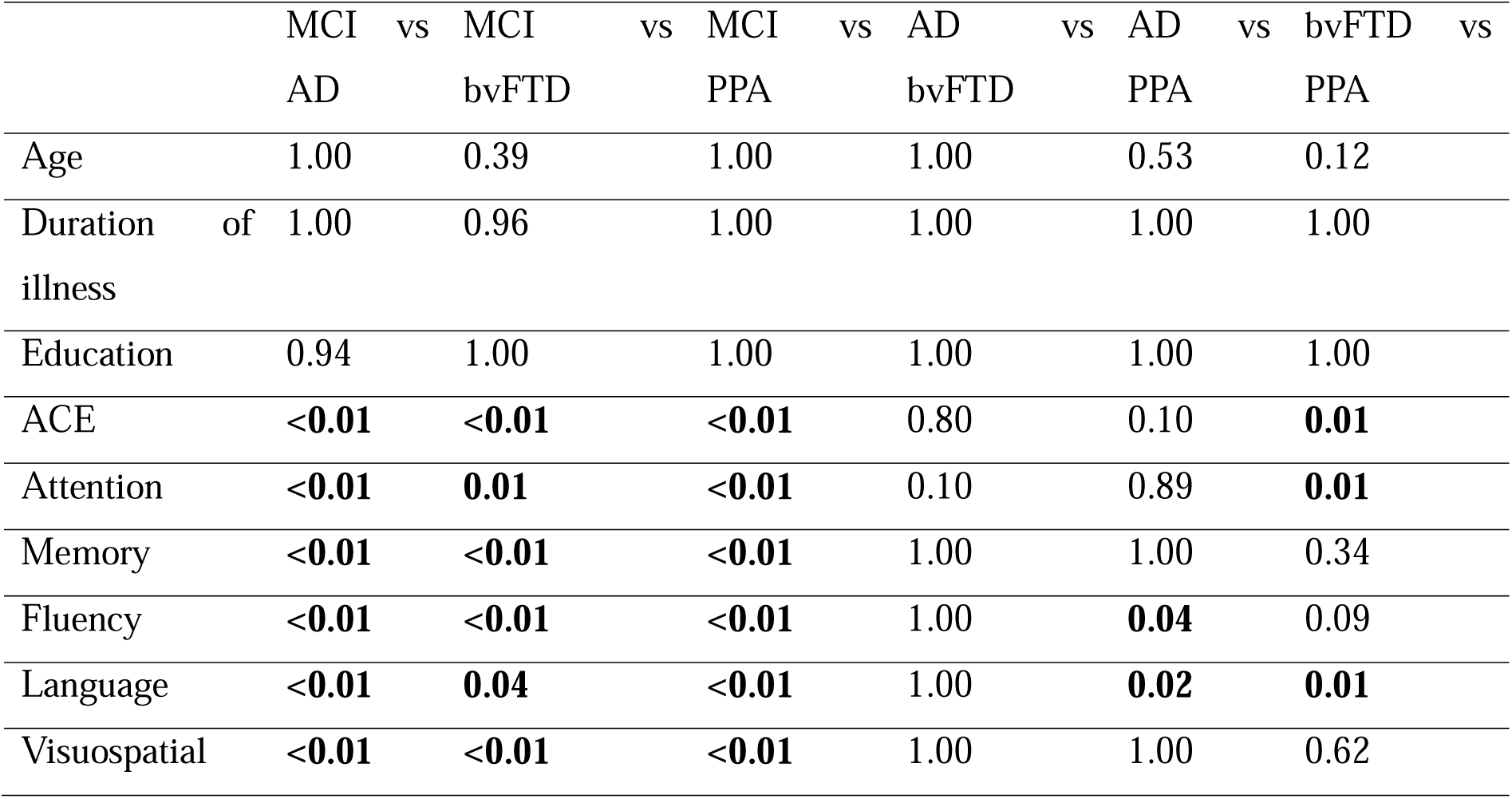
Multiple Comparisons (Post Hoc test-bonferroni).

